# Mining thermophile photosynthesis genes: a synthetic operon expressing *Chloroflexota* species reaction center genes in *Rhodobacter sphaeroides*

**DOI:** 10.1101/2025.09.22.677880

**Authors:** Yasir Rehman, Younghoon Kim, Michelle Tong, Ian K. Blaby, Crysten E. Blaby-Haas, J. Thomas Beatty

**Affiliations:** Department of Microbiology & Immunology, the University of British Columbia, Vancouver, BC V6T 1Z3, Canada; Department of Life Sciences, School of Science, University of Management and Technology, Lahore, Pakistan; U.S. Department of Energy Joint Genome Institute, Lawrence Berkeley National Laboratory, Berkeley, CA 94720, USA; Environmental Genomics and Systems Biology Division, Lawrence Berkeley National Laboratory, Berkeley, CA, 94720, USA; The Molecular Foundry, Lawrence Berkeley National Laboratory, Berkeley, CA 94720, USA

**Keywords:** *Chloroflexota*, photosynthetic reaction center, heterologous expression, anoxygenic, hot springs, bioinformatic database

## Abstract

Photosynthesis is the foundation of the vast majority of life systems, and therefore the most important bioenergetic process on earth, and the greatest diversity in photosynthetic systems are found in microorganisms. However, understanding of the biophysical and biochemical processes that transduce light to chemical energy has derived from the relatively small subset of proteins from microbes that are amenable to cultivation, in contrast to the huge number of microbial DNA sequences encoding proteins that catalyze the initial photochemical reactions that has been deposited in databases, such as from metagenomics. We describe the use of a *Rhodobacter sphaeroides* laboratory strain for expression of heterologous photosynthesis genes to demonstrate the feasibility of mining this resource, focusing on hot spring *Chloroflexota* gene sequences. Using a synthetic operon of genes, we produced a photochemically active complex of reaction center proteins in our biological system. We also present bioinformatic analyses of anoxygenic type II reaction center sequences from metagenomic samples collected from hot (42-90° C) springs available through the JGI IMG database, to generate a resource of diverse sequences that potentially are adapted to photosynthesis at such temperatures. These data provide a view into the natural diversity of anoxygenic photosynthesis, through a lens focused on high-temperature environments. The approach we took to express such genes can be applied for potential biotechnology purposes as well as for studies of fundamental catalytic properties of these heretofore inaccessible protein complexes.

## 1 Introduction

Prokaryotes in the phylum *Chloroflexota* (formerly *Chloroflexi*) are metabolically very diverse, and have been found in abundance in extreme environments ranging from freshwater hot springs to the sediments of the sea floor [3]. *Chloroflexota* species have been used as sources of enzymes with unusual catalytic properties and tolerance of temperature or pH extremes, for example, for biotechnology applications [4].

Most photosynthetic *Chloroflexota* contain a type II reaction center (RC), exemplified by the thermophiles *Chloroflexus aurantiacus* and *Roseiflexus canstenholzii*, and perform an-oxygenic photosynthesis utilizing an RC that contains bacteriochlorophyll (BChl) *a* and resembles the widely conserved RC of purple phototrophic bacteria such as *Rhodobacter* species [5,6]. In fact, all bacterial type II RCs are structurally very similar in terms of the two chlorophyll-binding proteins (PufL and PufM) that are universally present. The RC is imbedded in the cell membrane by virtue of transmembrane segments of PufL and PufM -- typically five alpha helices in each. The purple bacterial RC also contains a third protein, the H protein, which is absent from *Chloroflexota* RCs. The H protein stabilizes the RC, and it is unknown how the *C. aurantiacus* and *R. castenholzii* RCs are inherently resistant to the relatively high temperatures of environments in which these organisms are found. As shown in Figure 1, the cofactors present in type II RCs consist of six molecules of BChl and bacteriopheophytin (BPhe), two quinones (Q_A_ and Q_B_) and an iron atom. The 3-D organization of these cofactors is very similar in diverse species, although the ratio of BPhe:BChl may vary from 2:4 to 3:3, and the quinones may be solely ubiquinone or menaquinone, or one of each, depending on the species [7]. The *C. aurantiacus* and *R. castenholzii* RCs contain 3 BPhes and 3 BChls, and menaquinone is the sole type of quinone.

**Figure 1.**
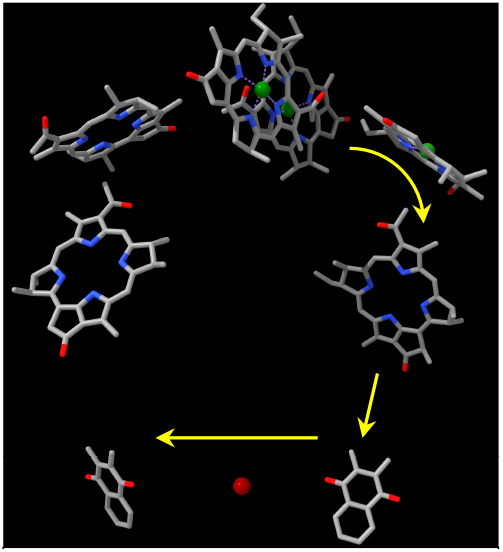
Spatial organization of RC cofactors. Molecules have been simplified and phytyl tails removed for clarity. Green spheres represent Mg^2+^ and the red sphere Fe^2+^. Top center, BChl dimer, P; top left, Bhe; top right, BChl; center left and right, BPhe; bottom right, menaquinone QA; bottom center, iron atom; bottom left, menaquinone Q_B_. Arrows show the pathway of electron transfer. Image is based on the *R. castenholzii* structure 5YQ7.pdb [1]. Image created using Chimerax [2].

Light energy excites a special pair of BChls (called P) in the RC, and electrons are transferred from P to a quinone (Q_B_) that leaves the RC after being reduced by acquisition of two electrons and two protons. The reduced quinone diffuses to a cytochrome complex where it is oxidized, while an oxidized quinone takes its place in the RC for another cycle of light-driven electron and proton transfer reactions.

The *C. aurantiacus* RC was isolated in 1993 [8], and since that time has been studied biochemically and spectroscopically. Electron microscopy has shown that the RC is surrounded by a ring of light harvesting proteins [9] that together are called the core complex, similar to the high resolution cryo-EM structures of core complexes from many species of purple phototrophic bacteria and *R. castenholzii* [10]. Although the *R. castenholzii* RC-LH core 3-D structure is known at a high resolution [1], and a number of spectroscopic tools have been used to study the function of this and the *C. aurantiacus* core complex [11-14] there is no genetic system for expressing mutant or engineered genes of these simple, thermostable RCs. There are basic scientific questions, such as the reasons for thermal stability, that could be addressed using a genetic approach. There are also potential bio-technical applications for thermostable RCs that harvest solar energy for electrical power, which would be facilitated by the use of genetic techniques. Furthermore, the number of *Chloroflexota* and related photosynthesis gene sequences is burgeoning and has far outstripped the number of organisms that have been cultivated. If these genes could be expressed in a tractable host that provided an appropriate membrane system, BChl, and other cofactors needed for assembly of a functional RC, it would greatly expand the diversity of such photosynthetic complexes that could be used to address fundamental scientific questions and exploited in biotechnology applications.

A major obstacle to the use of *E. coli* or other chemotrophic organism as a host for heterologous expression of photosynthesis genes is that it is not yet possible to obtain a high level of chlorophyll production in such species [15]. We circumvented this obstacle by using an engineered strain of the purple phototrophic bacterium *Rhodobacter sphaeroides* to express the *Chloroflexota* bacterium *L*.*E*.*CH*.*39_1* RC gene sequences found in a hot spring metagenome [16]. We show that these previously uncharacterized *Chloroflexota* genes were expressed in the *R. sphaeroides* host, using a synthetic, codon-optimized operon, resulting in an RC with an absorption spectrum that resembles that of *C. aurantiacus* and *R. castenholzii*. This RC was catalytically functional on the basis of light flash-induced oxidation of the P BChls. These results show that this system for heterologous expression of photosynthesis genes from uncultivated organisms has the potential for greatly expanding the diversity of naturally evolved RCs that can be studied, as material for future scientific investigations and biotechnological applications.

As such, we also searched the Joint Genome Institute’s (JGI) Integrated Microbial Genomes and Microbiomes (IMG/M) database for metagenomic-derived sequences encoding homologues of the RC proteins PufM and PufL that were obtained from aquatic environments in the temperature range of 42-90° C. In addition to identifying putative RC proteins from yet-to-be isolated hot spring bacteria, these comprehensive sequence analyses highlight the existence of highly divergent clades of PufL/PufM proteins encoded by *Chloroflexota* genomes and related environmental samples. Evolutionarily distinct hot spring metagenomic-derived RC proteins more closely related to proteins from Pseudomonadota were also found and represented the largest number of unique RC protein sequences identified in this analysis. In addition to providing a broad phylogenetic examination of the evolution of PufL/PufM, this analysis represents a resource that can be exploited to readily identify genes from organisms that are likely to be capable of performing anoxygenic photosynthesis in this temperature range.

## 2. Materials and Methods

### 2.1. Strains, growth conditions and plasmids

The *E. coli* strain DH5α was used for cloning, and strain S17-1 [17] was used for conjugation of plasmids into *R. sphaeroides. E. coli* BL21 DE3 [18] was used for isopropyl β-D1-thiogalactopyranoside (IPTG) induction of recombinant protein production for purification. *R. sphaeroides* strain RCx^R^ [19] containing plasmid pDJ-MK [20] was used as the host strain for plasmid expressing the *Chloroflexota* RC genes, and cultures were grown at 30° C in RLB medium [19] supplemented with tetracycline-HCl (0.5 µg/ml) and kanamycin sulfate (10 µg/ml).The *E. coli* plasmid-containing strain was grown at 37° C in Luria–Bertani (LB) medium [21] supplemented with antibiotics at the following concentrations in µg/ml: tetracycline, 10; kanamycin sulfate, 50.

Photosynthesis gene sequences of *Chloroflexota* bacterium *L.E.CH.39_1* [16] (NCBI accession number JACAEO010000223) encoding the RC were modified to match the codon usage of *Rhodobacter sphaeroides* and interspaced with ribosome binding site sequences designed for optimal expression as a synthetic operon [22,23]. Designed sequences were synthesized (Twist Bioscience, CA) and assembled using Gibson assembly into the pIND4 (linearized by *Nco*I + *Hind*III digest) expression plasmid containing an IPTG-inducible promoter and a kanamycin resistance marker [24], yielding plasmid pCFRC. A 6-His tag was incorporated immediately after the N-terminal methionine of the PufM protein. The DNA sequence of this operon is given in Supplementary Table S1, and a map of the recombinant plasmid pCFRC that expresses the *Chloroflexota* RC genes is shown in Supplementary Figure S1.

### 2.2. Protein purification, SDS-PAGE, and spectroscopy

The RC was purified as described previously [20], and summarized in the following text. Cells from induced cultures were collected by centrifugation, and disrupted in a French press cell. Membranes from disrupted cells were collected by ultracentrifugation, and solubilized in 1% *n*-dodecyl maltoside. Solubilized proteins were purified by binding to Ni^2+^-NTA agarose beads.

Proteins were separated in SDS-PAGE using 4% stacking and 12% resolving gels. Samples were mixed with 5x loading buffer and heated at the appropriate temperature for 10 minutes before loading. Gels were run at 150–200 V for ∼45 minutes or until the dye front reached the bottom of the resolving gel. For total protein visualization, gels were stained overnight with Coomassie brilliant blue and destained the following day.

For measurement of redox-induced changes in absorption, RC samples were reduced by the addition of an L-ascorbic acid solution to a final concentration of 10 mM, and oxidized by the addition of ferricyanide to 10 mM concentration. Photobleaching of RC samples was done using a 100 ms flash of illumination at 860 nm, while measuring the absorption at 865 nm as previously described [20].

### 2.3. Bioinformatic tools and data

Molecular graphics and analyses performed with UCSF ChimeraX, developed by the Resource for Biocomputing, Visualization, and Informatics at the University of California, San Francisco, with support from National Institutes of Health R01-GM129325 and the Office of Cyber Infrastructure and Computational Biology, National Institute of Allergy and Infectious Diseases.

Publicly available metagenomic bins, metagenomic-derived protein sequences, and associated search tools were accessed through the Joint Genome Institute’s IMG/MER (the login version of IMG/M) (April 2025) *[25]*. The UniProt database (version: 2025_01) *[26]* was accessed through the EFI-EST tool *[27,28]* using the Protein Family Addition Option with PF00124; to reduce the number of nodes, the UniRef90 *[29]* option was selected. Sequence Similarity Networks (SSNs) were visualized in Cytoscape (version 3.10.3) with the Prefuse Force Directed OpenCL layout without edge weights *[30]*. InterProScan *[31]* was used to determine the sequence coordinates of PF00124 domains *[32]*. Phylogenetic trees were built using MAFFT *[33]* for multiple sequence alignments and FastTreeMP *[34]* with default parameters on the CIPRES Science Gateway *[35]*. The resulting trees were visualized and annotated in iTOL *[36]*. Multiple sequence alignments were manually edited in Jalview *[37]* to remove fragments and poorly aligned regions; sequence redundancy in the edited alignment was reduced based on 95% sequence identity using the Remove Redundancy tool in Jalview, *i.e*., if the percentage identity between the aligned positions of any two sequences exceeds 95%, the shorter sequence is discarded. Protein sequences that remained after this filtering step are referred to as “unique”. Gene neighborhoods for selected proteins from UniProt were visualized with EFI-GNT *[28]*. Data and information for SSNs, the alignment, the tree, and retrieved gene neighborhood information are available in Tables S2, S3, and S4.

#### 2.2.1. Curation of hot spring metagenomes

Identification of hot spring metagenomic bins involved a search of public IMG metadata for user-supplied sample collection temperature of above 42° C associated with datasets labeled as Thermal springs: Hot (42-90° C). Not all samples in this Ecosystem Subtype were collected from waters in that temperature range, and those samples were excluded in this analysis. For missing metadata, we curated sample collection temperatures from the associated published studies, resulting in a conservative set of metagenomics-derived sequences isolated from samples of at least 42° C. This approach reduced the number of metagenomic bins from 3,641 to 233 with a confirmed collection temperature between 42° C and 90° C (Table S5). These 233 metagenomic samples are referred to as “hot spring metagenomic bins”, and proteins from these bins as “hot spring proteins”. The resulting curated data are associated with the studies of [38-44]. For UniProt- or NCBI-derived protein sequences, we collected isolation source from the corresponding GenBank record.

#### 2.2.2. Identification of reaction center proteins PufL and PufM

We searched the predicted protein sequences derived from the hot spring metagenomic bins using PF00124 and a blastp search with PufL and PufM from *Chloroflexota* bacterium L.E.CH.39_1 and the fused PufL-PufM protein from *Roseiflexus* sp. 629F.7. The blastp search was used to identify homologous proteins that do not have a significant match to PF00124, which can be true for related but highly divergent sequences. The blastp search identified 9 sequences from hot spring metagenomic bins the that are not recognized as belonging to PF00124, for a total of 29,685 protein sequences. An additional 51,363 UniProt sequences were identified using PF00124. Neither the blastp search nor the PF00124 search distinguishes PufM and PufL from the homologous D1 (PsbA) and D2 (PsbD) photosystem II (PSII) reaction center proteins from cyanobacteria, algae and land plants. To distinguish PufL and PufM from PsbA and PsbD, we generated an SSN with an alignment score of 90, which clearly delineates clusters containing PsbA or PsbD from clusters containing PufL or PufM (Figure S2).

To further analyze PufL and PufM, additional sequence analyses was performed. For each node in a Puf cluster (Figure S2), the member with the longest sequence was kept for downstream analysis. PufL and PufM are predicted to be encoded by a single ORF in some *Chloroflexota* genomes [45,46]. There is evidence that the mature form of these gene products is separate PufL and PufM proteins [1], but it is unknown at what point the fused polypeptide is processed or whether the RNA is processed. Nevertheless, a PufL-PufM fusion protein is computationally predicted during genome annotation in these cases. Therefore, we used InterProScan and alignments to PF00124 to identify the sequence coordinates for individual domains in both single- and double-domain proteins.

This resulted in the identification of 164 PufL-PufM fusions from hot spring metagenomic bins and 20 representative fusions from UniProt. In Tables S2 and S3, domains from a fusion protein are labeled with “N” or “C”. The sequences corresponding to each domain (either from a fusion or non-fusion protein) were then used to regenerate the SSN, using alignment scores of 50 or 80. Lower alignment scores result in more edges connecting nodes, while higher alignment scores result in less edges. Use of multiple alignment scores allows the visualization of similarity between representative protein sequences at different thresholds. For each network, a node represents a cluster of sequences that share 95% sequence identity calculated by EFI-EST. Representative sequences and cluster members can be found in Table S2. Representative sequences from each node in the SSN generated with an alignment score of 80 were used to generate a multiple sequence alignment and phylogenetic tree (described in more detail in previous section). PsbA1, PsbA2, and PsbD1 from *Acaryochloris marina* were used to root the tree in iTOL.

Each subsequent analysis reduced the number of non-redundant protein sequences. Protein sequences were considered redundant if they shared 95% sequence identity or higher to one another. With metagenomic-derived proteins, using sequence identity to cluster sequences can be potentially misleading, as two proteins would be considered to be less than 95% identical if one sequence is shorter by 5%. Therefore, the phylogenetic reconstruction represents a more conservative estimate of uniqueness, as the multiple sequence alignment was edited to remove short sequences and poorly aligned regions before reducing the number of representative sequences based on sequence identity. As a result, 1,716 protein sequences representing each connected node in the PufLM SSN was reduced to 995 representative sequences in the phylogenetic tree.

Following SSN construction, nodes without connecting edges were not retained in the subsequent analysis. In Table S2, these are listed as singletons under “cluster”. Typically, these nodes represent small incomplete protein sequences, *i.e*., fragments. However, the recently isolated PufL and PufM sequences in *Vulcanimicrobium alpinum* [47] were represented by edgeless nodes in the PF00124 SSN and were not included in downstream analyses. Therefore, to ensure that removal of node singletons did not inadvertently remove divergent full-length hot spring proteins, we performed a blastp search against the predicted proteins from each hot spring metagenomic bins using PufL from *V. alpinum*; we did not identify any similar hot spring proteins that are not already represented in the phylogenetic reconstruction.

## 3. Results

### 3.1. *Design, construction and expression of Chloroflexota synthetic* pufLM *operon encoding the RC*

The genome of *Chloroflexota* L.E.CH.39_1 (hereafter referred to as *Chloroflexota* sp.) currently is in 459 contigs [16], with the RC *pufL* and *pufM* genes adjacent to each other on one contig. The amino acid sequence identities in full-length (Needleman-Wunsch) alignments between the proteins encoded by *puf* gene homologues in *Chloroflexota* sp. and

*C. aurantiacus* are: *pufL*, 85%; *pufM*, 83%. For expression from the plasmid vector in the *R. sphaeroides* host, an operon (Figure 2) was synthesized with codons changed to the relatively high GC codons most frequently found in *R. sphaeroides* (see Materials and Methods section). An AlphaFold model [48] predicted the N-termini of PufL and PufM to be located in the cytoplasm (Figure 3). A 6x-His tag was incorporated at the N-terminus of the PufM protein to facilitate purification by metal ligand affinity chromatography. The operon was inserted into the IPTG-inducible expression plasmid pIND4, yielding plasmid pCFRC, which was introduced into the *R. sphaeroides* strain RCx^R^ (pDJ-MK) that lacks all *puf* genes encoding the RC and expresses menaquinone biosynthesis genes carried on a second plasmid [20].

**Figure 2.**
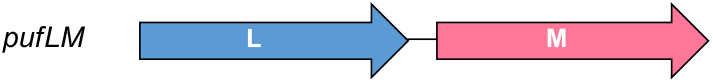
Representation of synthetic operon of *Chloroflexota* sp. RC genes pufL and pufM. Operon name is given on the left, arrows represent coding regions.

**Figure 3.**
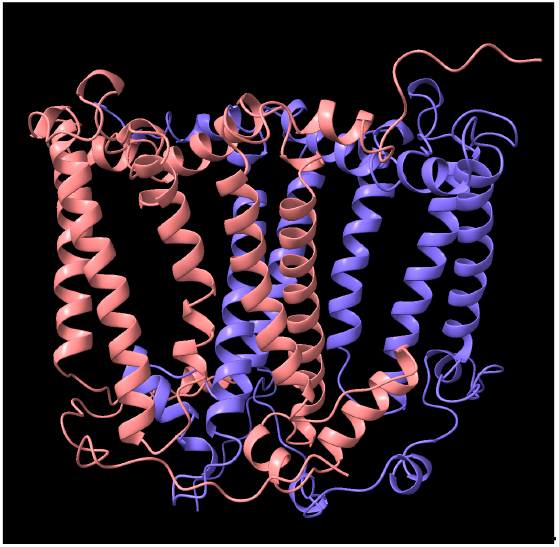
AlphaFold model of *Chloroflexota sp*. RC. PufL is in blue, PufM in pink. The periplasmic side is at the top, and the cytoplasmic side at the bottom, with the vertical alph-helices traversing the cytoplasmic membrane. Image obtained using Chimerax [2].

### 3.2 SDS-PAGE, absorption spectra and catalytic activity of the RC complex produced by the pufLM operon

The SDS-PAGE analysis of the RC proteins was complicated by their aberrant mobility and aggregation, which were affected by the temperature of sample treatment (Figure 4). Increasing the temperature of sample heating from 35 to 80° C prior to loading the gel resulted in a decrease in intensity of the faster-migrating bands, and the appearance of bands near the top of the gel. The masses of PufL and 6His-PufM are 35.1 and 35.5 kD; therefore, these proteins would migrate as a single band in the gel system employed. Furthermore, hydrophobic proteins such as PufL and PufM often migrate faster than would be indicated by their mass. Therefore, we suggest that the broad band at approximately the 27 kD position represents the PufL and 6His-PufM proteins, whereas the slower-migrating bands represent aggregations of these proteins.

**Figure 4.**
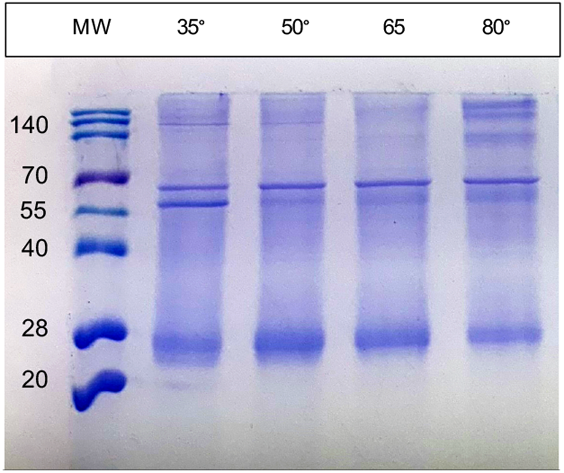
SDS-PAGE of *Chloroflexota sp*. RC showing effects of sample treatment temperature on protein migration. MW, molecular weight ladder; numbers above lanes give the temperature (celsius) of sample treatment prior to gel loading.

After affinity chromatography purification, the RC complex yielded absorption spectra with three peaks in the 700 to 800 nm range (the Qy bands), which is characteristic of anoxygenic Type II RCs (Figure 5). In this case, we attribute these peaks to the presence of the P BChls (∼860 nm), accessory BChl (∼810 nm), and BPhes (∼760 nm). In the visible (Qx) region of the spectrum the peak near 535 nm is attributed to BPhe, and the peak near 600 nm to BChl. The relative peak heights in both the Qy and Qx regions of the spectrum are consistent with the presence of three BChls and three BPhes per RC, as has been shown for the RCs of *C. aurantiacus* and *R. castenholzi* [8,46]. Figure 6 shows the ascorbate-reduced and ferricyanide-oxidized absorption spectra of the *Chloroflexota* sp. RC. The bleaching of the 860 nm peak after oxidation confirms the assignment of this band to the P BChls, and shows that they are potentially capable of initiating electron transfer to the RC quinones as diagrammed in Figure 1. To determine whether this *Chloroflexota* sp. RC is fully catalytically active (*i.e*., capable of photon-driven Q_B_ reduction as shown in Figure 1), we specifically excited the P BChls of the RC using a flash of 865 nm light, and followed the kinetics of bleaching and recovery of the 860 nm peak. The rate of recovery of the 860 nm peak reflects the rate of the back reaction of electron transfer from the quinones to P^+^, which differs depending on whether Q_B_ is present. As shown in Figure 7, there was a rapid decrease in absorbance at 860 nm after the flash, followed by a relatively slow recovery that started to plateau after about 5 s. These data show that this RC is fully functional in terms photon-driven electron transfer from the P BChls to the second of the two quinones, Q_B_. This interpretation is based on the rate of recovery, which in the *R. sphaeroides* RC takes about 1 s from Q_B_^-^, whereas recombination from Q_A_^-^ takes only about 0.1 s [49]. The *R. castenholzii* and *C. aurantiacus* RCs similarly have fast recombination rates from Q_A_^-^, on the order of 0.04 to 0.06 s [11,50]. Therefore, the kinetics of recovery that we observed for this *Chloroflexota* RC are closer to the values for recovery by electron transfer to P^+^ from Q_B_^-^ than from Q_A_^-^ in homologous RCs, and indicates that this RC is replete in cofactors and fully functional.

**Figure 5.**
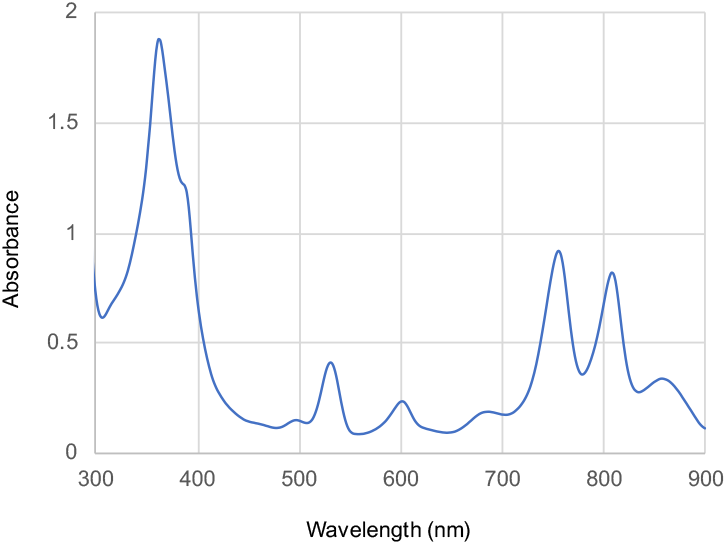
Absorption spectrum of the *Chloroflexota* sp. RC.

**Figure 6.**
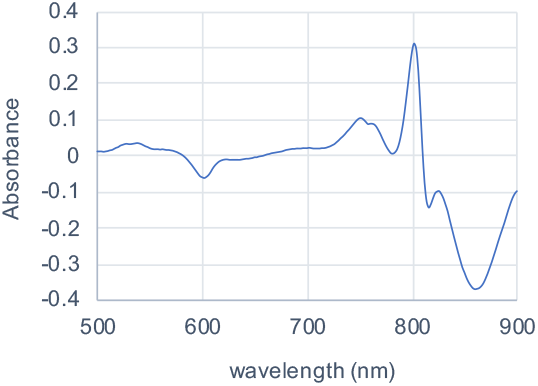
Oxidized minus reduced *Chloroflexota* sp. RC absorbance spectrum.

**Figure 7.**
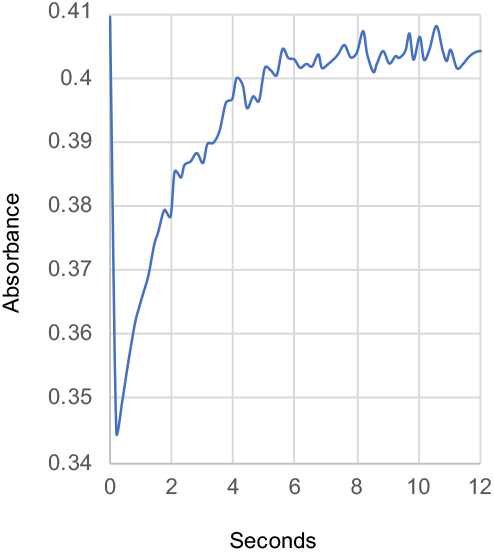
Kinetics of recovery of *Chloroflexota* sp. RC absorbance at 860 nm after a flash of 865 nm illumination.

### 3.4. Bioinformatic analyses of RC proteins

Given the success with heterologous expression of metagenomic-derived *Chloroflexota* sp. proteins to study functions encoded by non-cultivable bacteria, we next aimed to generate a resource of diverse metagenomic sequences potentially adapted to anoxygenic photosynthesis at extreme temperatures. We searched publicly available metagenomic samples collected from hot (42-90° C) springs available through the JGI IMG database and focused on homologs of PufL and PufM. Identified sequences were clustered to reduce the number of identical and highly similar sequences and were compared to homologous proteins available in the UniProt database, which were also clustered to reduce the number of identical and highly similar sequence. Based on an iterative approach that combines SSNs (Figure 8 and Figure S2) with phylogenetic reconstruction (Figure 9), we identified 386 unique PufL homologs, 577 unique PufM homologs, 14 N-terminal unique PufL-domains, and 18 C-terminal unique PufM-domains. (See Materials and Methods for a discussion on identification of proteins and defining uniqueness.) These homologs and related sequences are grouped into 8 clusters in the network and 8 corresponding clades in the tree; 4 for PufL and 4 for PufM (Figure 8 and Figure 9). The N- and C-terminal domains were predicted from genes encoding PufL-PufM fusions. There is evidence that the mature form of these gene products is separate PufL and PufM proteins [1], but it is unknown at what point the fused polypeptide is processed (or whether the RNA is processed). To avoid potential artifacts in the analyses, only sequence corresponding to the domain PF00124 was used for comparisons, and “N_” is used to distinguish the PufL-like region from these fusions and “C_” is used to distinguish the PufM-like region.

**Figure 8.**
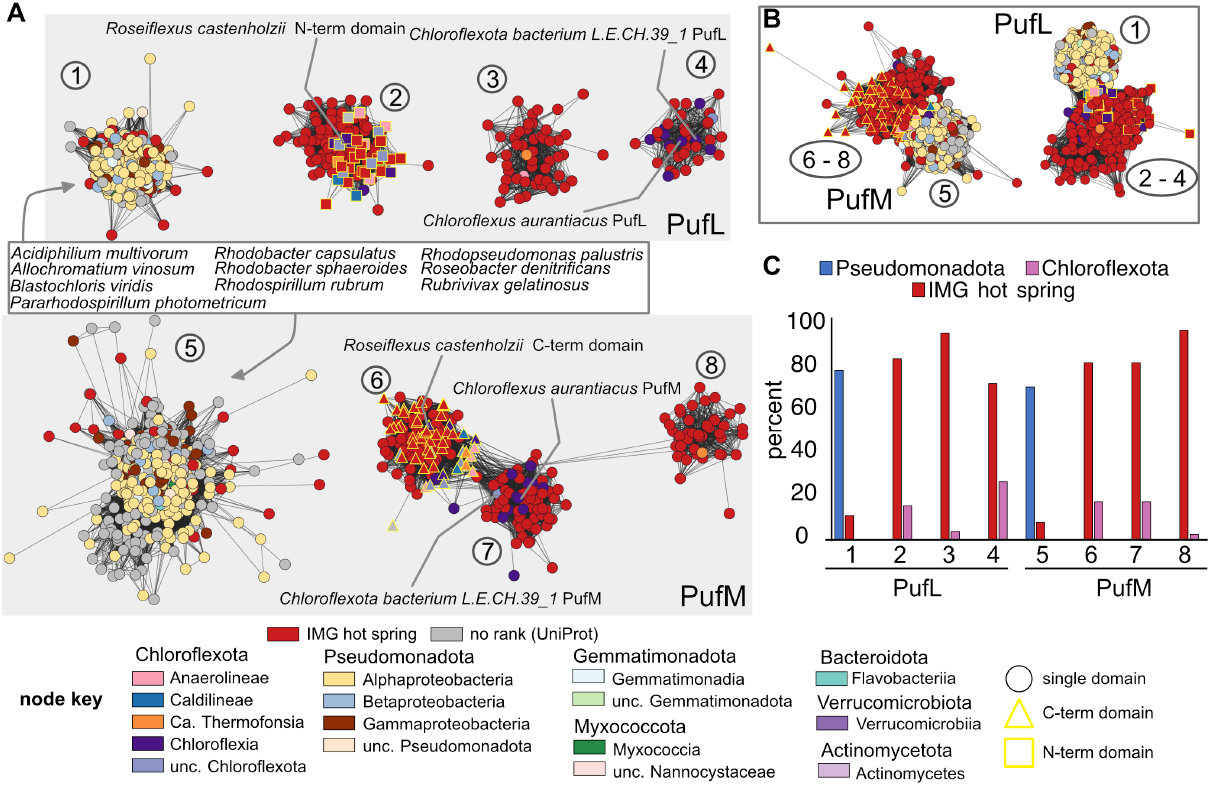
Identification of PufL and PufM proteins encoded by metagenomic samples from hot springs. **A**, sequence similarity network of PufL-like and PufM-like proteins using an alignment score of 80. Clusters 1 – 4 represent PufL-like proteins based on sequence similarity to characterized PufL sequences; clusters 5 through 8 represent PufM-like proteins based on sequence similarity to characterized PufM sequences. Nodes are colored by class according to the node key, colored red for hot spring proteins identified in IMG, or colored grey for proteins from UniProt with no assigned taxonomic rank. Square nodes represent N-terminal PufL-like domains, and triangles are used to indicate those nodes representing C-terminal PufM-like domains from a fusion protein. **B**, the same network as in panel **A**, except that an alignment score of 50 was used to draw edges. Nodes are colored the same as in panel **A. C**, number of nodes represented by the indicated taxa as a percent for each cluster. Minor taxa are not shown. A graphical representation of all taxa can be found in Figure S3.

**Figure 9.**
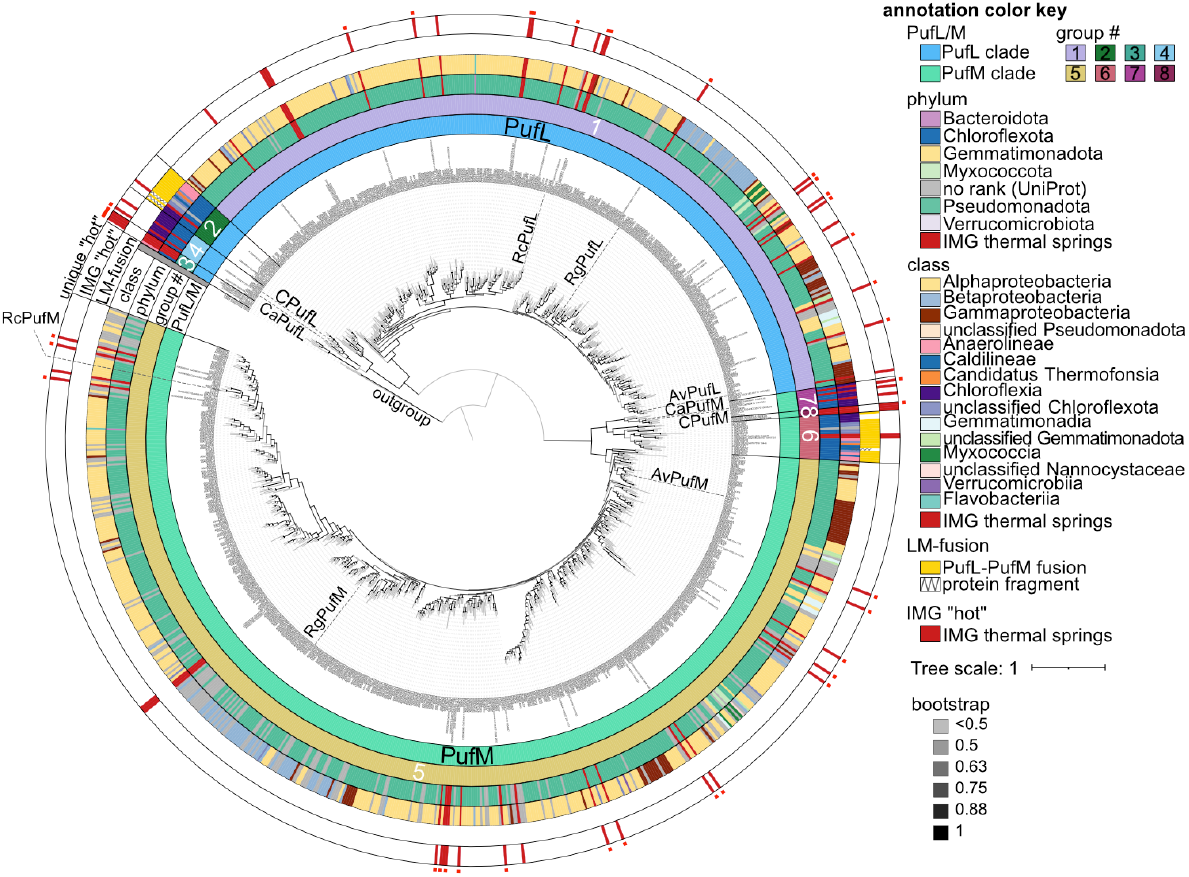
Phylogenetic reconstruction of PufL and PufM homologs. Outer ring annotations are colored according to the color key. Leaves are labeled that represent PufL and PufM from *Chloroflexota* bacterium L.E.CH.39_1 (C) or characterized proteins in SwissProt. Av, *Allochromatium vinosum*; Ca, *Chloroflexus aurantiacus*; Rc, *Rhodobacter capsulatus*; Rg, *Rubrivivax gelatinosus*. To help identify the leaves representing IMG thermal spring sequences, red bars are used for phylum, class, and in the IMG “hot” annotation ring. IMG hot spring proteins with less than 95% identity to another protein in Genbank are labelled with a small red square on the outmost of the circle. PufL-PufM fusions were found exclusively in the clades representing cluster 2 (contains the PufL-like domain) and cluster 6 (contains the PufM-like domain). In those cases where two domains were not identified for proteins in these clusters, some of those proteins are fragments (annotated with a zigzag line), the sequence encoding PufM is at the 5’ most end of the contig (A0A0P9DKP2), or the gene model appears to be incorrect (A0A7W1TQK3), suggesting that the full fusion was not captured in these cases. The tree was rooted at the outgroup (PsbA1, PsbA2, and PsbD1 from *Acaryochloris marina*).

Based on a sub-sampled gene neighborhood analysis, photosynthesis operons typically contain genes for both PufL and PufM (Figure 10 and Table S2). This trend holds true even for sequences from clusters PufL_4 and PufM_7 where the photosynthesis genes are separated into two distinct genomic locations, one containing adjacent genes for PufL and PufM and the second locus containing adjacent genes for PufA, PufB and PufC (Figure 10). Therefore, the unequal number of non-redundant PufL and PufM proteins is likely due to a combination of a small bias in the number of PufM homologs initially identified (∼25% more PufM than PufL) and a larger bias in the similarity among PufM homologs compared to among PufL homologs. As seen in Figure 8A, the main PufM cluster (cluster 5) is less compact than the main PufL cluster (cluster 1), which reflects a larger proportion of low alignment scores between PufM homologs, *i.e*., fewer edges between PufM nodes. The extent of divergence between PufM homologs compared to PufL homologs is also apparent in the phylogenetic tree (Figure 9).

**Figure 10.**
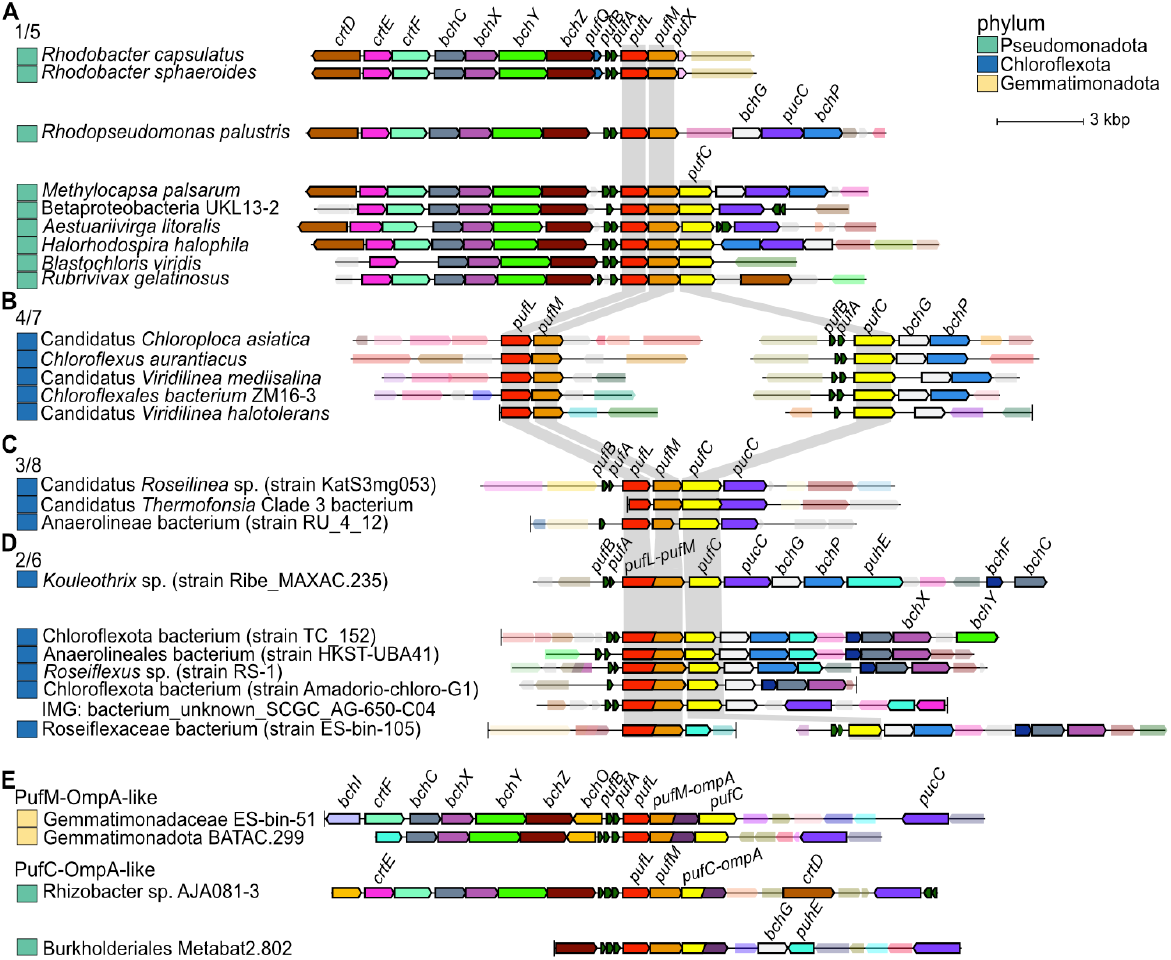
Gene neighborhoods containing *puf* genes. **A**, neighborhoods containing genes encoding proteins from PufL cluster 1 and PufM cluster 5. **B**, neighborhoods containing genes encoding proteins from PufL cluster 4 and PufM cluster 7. In these genomes, *pufC* is in a separate neighborhood. **C**, neighborhoods containing genes encoding proteins from PufL cluster 3 and PufM cluster 8. **D**, neighborhoods containing genes encoding fusions proteins that have domains from PufL cluster 2 and PufM cluster 6. **E**, examples of gene neighborhoods encoding PufM or PufC proteins with a C-terminal OmpA-like domain (dark purple). For all panels, genes encoding homologous proteins are colored the same, and unlabeled genes are transparent. A vertical black line is used to indicate the end of a contig. The corresponding phylum for each bacterium is given to the left, according to the color key. Grey background shading is used to emphasize the distinct arrangements of in each cluster of genes encoding PufL, PufM, and PufC.

As reflected in the SSN and phylogenetic tree, we identified 4 distinct clusters/clades for both PufL and PufM homologs (Figure 8 and Figure 9). The largest clusters (cluster PufL_1 and PufM_5) are dominated by sequences from *Pseudomonadota*, while the other clusters (PufL_2, PufL_3, PufL_4, PufM_6, PufM_7, and PufM_8) are dominated by sequences from *Chloroflexota* and the hot spring samples (Figure 8B). The N-terminal PufL region and C-terminal PufM region from the fused gene products unique to *Chloroflexota* are found exclusively in clusters 2 and 6, respectively. We also identified three PufM homologs with a C-teminal OmpA-like domain in PufM cluster 5 (Figure 10E). The hypothesis that this OmpA-like domain may function in photosynthesis was further supported by the identification of analogous fusions where PufC homologs contain a OmpA-like N-terminal domain (Figure 10E). However, as these Puf-OmpA fusions are relatively rare, we can’t rule out the possibility of inaccurate gene models. In the case of genes encoding the fused PufL and PufM, a single ORF is a conserved characteristic for the proteins in cluster 2 and 6. Those sequences in either cluster 2 or 6 with only a single domain identified appear to be fragments.

Based on our iterative approach that combined SSNs and phylogenetic reconstruction, we identified 30 PufL sequences, 29 PufM sequences, and 2 C-terminal domains that are unique and represented by an IMG hot spring sequence (Table S6). As the UniProt database is not an exhaustive collection of predicted proteins, we searched NCBI’s Genbank using a threshold of 95% identity to determine if similar proteins have been detected in deposited sequencing projects. This analysis suggests 15 unique hot spring sequences in PufL_1, 3 in PufL_3, 1 in PufL_4, 17 in PufM_5, 1 in PufM_7, and 1 in PufM_8 (Figure 9 and Table S6). The closest homologs of these proteins are also often from hot spring samples. Exceptions do occur, such as three Puf sequences from IMG identified from the Seven Mile Hole Area in Yellowstone National Park with closest matches (83-85% identity) to a *Pseudomonadota* bacterium sequenced from plastic particles (plastisphere) in the Pacific Ocean, and two IMG Puf sequences from Empress Pool OF2, Yellowstone National Park, with closest matches (88% identity) to sequences from plastic particles in the Atlantic Ocean [51].

## 4. Discussion

In this paper we describe how the genetically tractable alphaproteobacterium *R. sphaeroides* was used as a chassis for the expression of distantly related *Chloroflexota* photosynthesis genes mined from hot spring metagenomic samples. Although *Chloroflexota* sp. has never been cultivated or microscopically visualized, the RC proteins are 82-85% identical in sequence alignments to those of *C. aurantiacus*, whereas the corresponding identities between those of *R. castenholzii* and the *Chloroflexota* sp. are 49% (PufL) and 48% (PufM). These relative similarities are implicit in the SSNs and phylogenetic tree generated in our bioinformatic analyses. Therefore, the photosynthetic apparatus of *Chloroflexota* sp. appears to be physiologically similar to that of *C. aurantiacus*, although there is no evidence of chlorosome-related proteins encoded in the contigs. Attempts to obtain anaerobic phototrophic growth of *R. sphaeroides* strain RCx^R^ [19] containing plasmids pDJ-MK [20] and pCFRC were unsuccessful, which could be due to an inability of the *R. sphaeroides* cytochrome *bc1* complex, which oxidizes ubiquinol [5], to oxidize menaquinol produced by the *Chloroflexota* sp. RC.

Our bioinformatic analysis of PufL and PufM sequences shows that hot spring sequences almost exclusively cluster with sequences from Chloroflexota, supporting previous analyses that Chloroflexota are abundant in hot spring microbial samples [52]. Not only are they abundant, but the Chloroflexota proteins can be grouped into one of three ancient lineages supported by the sequence analyses and distinct operonic structure of photosynthesis genes. Intriguingly, a conservative approach that takes into account the sequence identity of aligned positions in the multiple sequence alignment suggests that a greater number of unique hot spring sequences (*i.e*., less than 95% identity to proteins deposited in UniProt) are related to proteins from *Pseudomonadota*. As such, even though *Chloroflexota* PufL and PufM sequences dominate the hot spring samples and are derived from distinct and ancient lineages, *Pseudomonadota*-like metagenomic-derived sequences represent a broader swath of sequence diversity.

The overwhelming number of diverse and unique PufL and PufM sequences have yet-to-be experimentally characterized. This lack of data is a combination of limited availability of genetically tractable model systems, the inability to cultivate a large number of anoxygenic phototrophs detected with DNA sequencing, and rapidly growing collections of predicted protein sequences that are impractical for experimental characterization. Here, we present a resource of unique and diverse PufL and PufM sequences for selecting candidate sequences, and an experimental approach to study these proteins through heterologous expression. Combined with the decreasing cost of DNA synthesis, these resources can enable the study and genetic modification of novel photosynthesis proteins from thermophiles that may never be amenable to cultivation as pure cultures in the laboratory.

## Supporting information

compressed supplementary files

## Supplementary Materials

The following supporting information can be downloaded at: www.mdpi.com/xxx/sx: Figures S1-S3; Tables S1-S6.

## Author Contributions

Conceptualization, J.T.B., I.K.B. and C.E.B-H.; methodology, J.T.B., I.K.B. and C.E.B-H.; resources, J.T.B., I.K.B. and C.E.B-H; data curation, C.E.B-H.; writing—original draft preparation, J.T.B. and C.E.B-H.; writing—review and editing, Y.R., Y.K., M.T., I.K.B.; supervision, J.T.B. and I.K.B.; funding acquisition, J.T.B. and C.E.B-H. All authors have read and agreed to the published version of the manuscript.

## Acknowledgments

Molecular graphics images were prepared using UCSF ChimeraX [[UCSF ChimeraX: Tools for structure building and analysis](https://onlinelibrary.wiley.com/doi/10.1002/pro.4792). Meng EC, Goddard TD, Pettersen EF, Couch GS, Pearson ZJ, Morris JH, Ferrin TE. Protein Sci. 2023 Nov;32(11):e4792], developed by the Resource for Biocomputing, Visualization, and Informatics at the University of California, San Francisco, with support from National Institutes of Health R01-GM129325 and the Office of Cyber Infrastructure and Computational Biology, National Institute of Allergy and Infectious Diseases.

DNA synthesis and expression constructs were provided by the U.S. Department of Energy Joint Genome Institute (https://ror.org/04xm1d337), a DOE Office of Science User Facility, supported by the Office of Science of the U.S. Department of Energy operated under Contract No. DE-AC02-05CH11231 (proposal:510024, doi.org/10.46936/10.25585/60008867).

## Abbreviations

The following abbreviations are used in this manuscript:

RC(s): Reaction center(s)
BChl(s): Bacteriochlorophyll(s)
BPhe(s): Bacteriopheophytin(s)
P: Primary donor or special pair of BChls in the RC
Q_A_: Quinone within the RC
Q_B_: Quinone within the RC
sp.: Species
ms: millisecond
nm: nanometer
NTA: nitrilotriacetic acid
JGI: Joint Genome Institute
IMG/M: Integrated Microbial Genomes and Microbiomes
SSN(s): Sequence similarity network(s)

