## Supplementary material for "Mining thermophile photosynthesis genes: a synthetic operon expressing *Chloroflexota* species reaction center genes in *Rhodobacter sphaeroides*": compressed supplementary files: Rehman et al. supplementary figures S1-S3.pdf

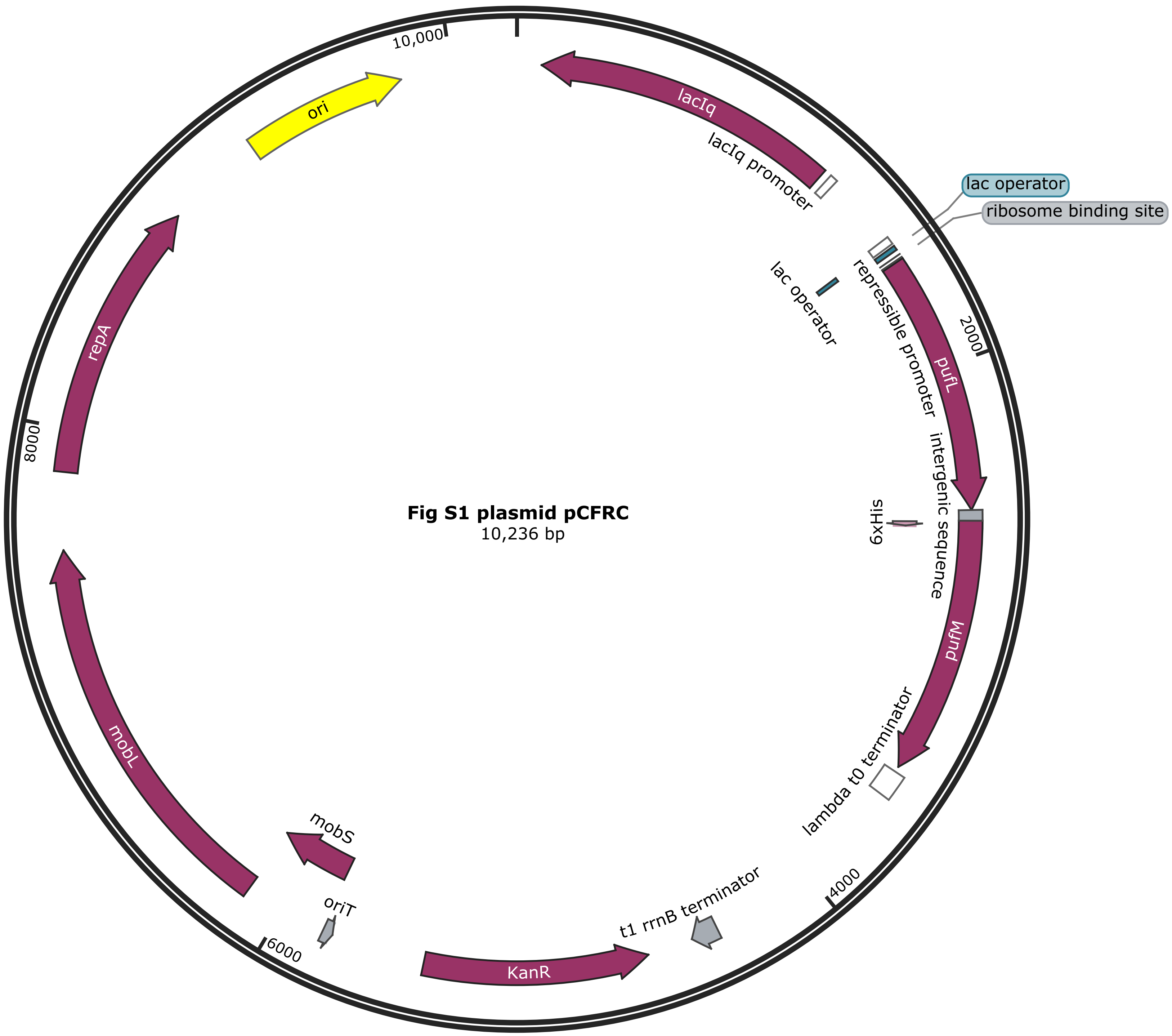
Created with SnapGene®
Fig S1 plasmid pCFRC
10,236 bp
10,000
2000
4000
6000
8000
lacIq
lacIq promoter
lac operator
ribosome binding site
lac operator
repressible promoter
pufL
intergenic sequence
6xHis
pufM
lambda t0 terminator
t1 rrnB terminator
KanR
oriT
mobS
mobL
repA
ori

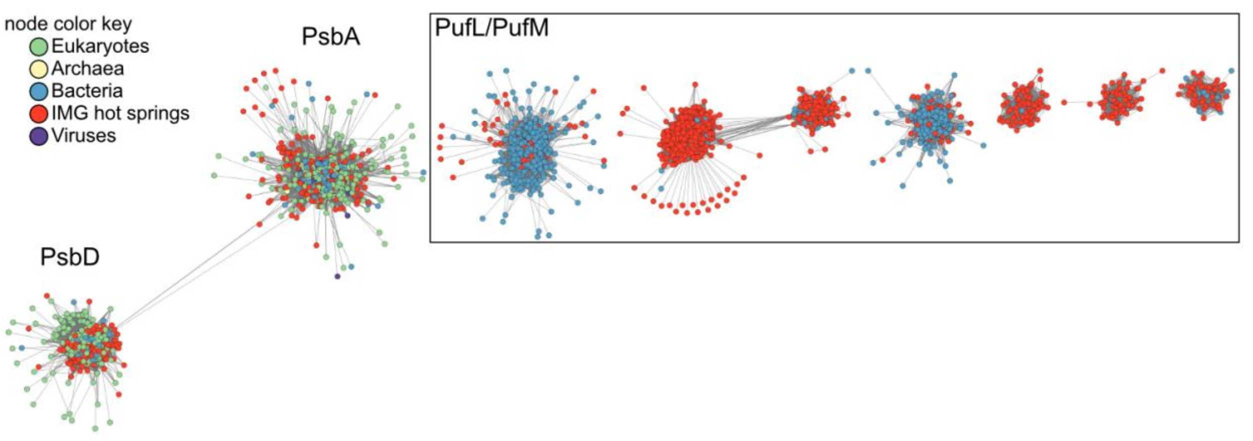


Figure S2. Identification of PufL and PufM proteins encoded by metagenomic samples from hot springs. Sequence similarity network of proteins with matches to PF00124. Nodes are colored according to the node key.

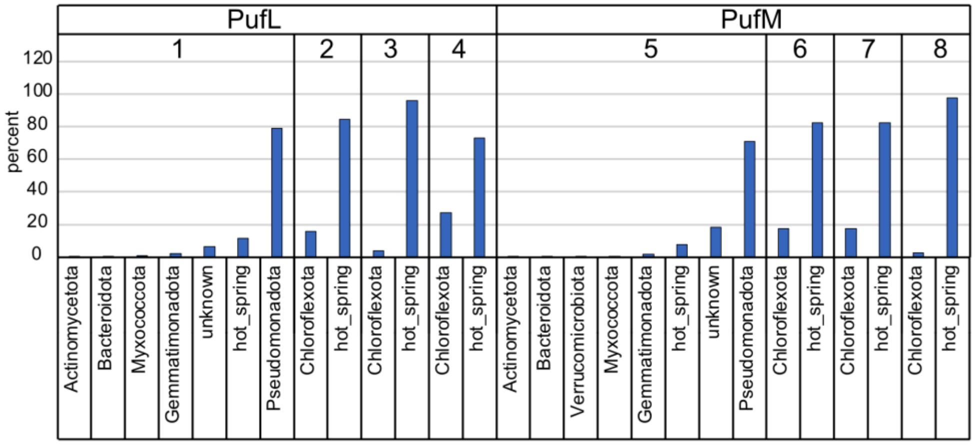


Figure S3: Representative taxa for each node in Figure 8. For each node in the PufL-PufM SSN, the number of known taxa were calculated from the node members. The most abundant known phylum was used as the representative taxon. The percentage abundance was calculated by dividing the number of nodes with a given representative taxon by the total number of nodes in a given cluster.
